# Representation of contralateral visual space in the human hippocampus

**DOI:** 10.1101/2020.07.30.228361

**Authors:** Edward H Silson, Peter Zeidman, Tomas Knapen, Chris I Baker

## Abstract

The initial encoding of visual information primarily from the contralateral visual field is a fundamental organizing principle of the primate visual system. Recently, the presence of such retinotopic sensitivity has been shown to extend well beyond early visual cortex to regions not historically considered retinotopically sensitive. In particular, human scene-selective regions in parahippocampal and medial parietal cortex exhibit prominent biases for the contralateral visual field. Here we used fMRI to test the hypothesis that the human hippocampus, which is thought to be anatomically connected with these scene-selective regions, would also exhibit a biased representation of contralateral visual space. First, population receptive field mapping with scene stimuli revealed strong biases for the contralateral visual field in bilateral hippocampus. Second, the distribution of retinotopic sensitivity suggested a more prominent representation in anterior medial portions of the hippocampus. Finally, the contralateral bias was confirmed in independent data taken from the Human Connectome Project initiative. The presence of contralateral biases in the hippocampus – a structure considered by many as the apex of the visual hierarchy - highlights the truly pervasive influence of retinotopy. Moreover, this finding has important implications for understanding how this information relates to the allocentric global spatial representations known to be encoded therein.

**Significance Statement:** Retinotopic encoding of visual information is an organizing principle of visual cortex. Recent work demonstrates this sensitivity in structures far beyond early visual cortex, including those anatomically connected to the hippocampus. Here, using population receptive field modelling in two independent sets of data we demonstrate a consistent bias for the contralateral visual field in bilateral hippocampus. Such a bias highlights the truly pervasive influence of retinotopy, with important implications for understanding how the presence of retinotopy relates to more allocentric spatial representations.

## Introduction

The segregation of visual information processing from the two visual fields, with biased representation of the contralateral visual field, is a fundamental feature of the human visual system (Wandell et al., 2007). Although historically considered a feature reserved for the earliest stages of visual cortex (V1-V4), recent work highlights privileged processing of contralateral space throughout the brain (Kravitz et al., 2013; Silson et al., 2015). Indeed, at least twenty separate maps of the visual field have been identified throughout cortex (Wandell et al., 2007; Swisher et al., 2015) and greater contralateral sensitivity has been reported in anterior regions of ventral temporal cortex (Hemond et al., 2007; Kravitz et al., 2010; Chan et al., 2010) and the default mode network (Szinte & Knapen, 2020). Further, retinotopic maps of the contralateral visual field have been reported in the frontal eye fields (Mackey et al., 2017), the frontal lobes (Silver & Kastner, 2009) and even the cerebellum (Van Es et al., 2019). Given the seemingly ubiquitous influence of contralateral visual encoding, we asked whether the human hippocampus – a structure critical for long-term episodic memory (Scoville & Milner, 1957; Squire, 1992) and spatial navigation (O’Keefe & Nadal, 1978) among many other cognitive functions – also exhibits a contralateral bias for visual space.

Although at first glance, the notion of retinotopic sensitivity within the hippocampus may seem surprising, there is growing evidence to suggest that such sensitivity may nonetheless exist.

For example, the hierarchical model of visual processing proposed by Felleman and Van Essen places the hippocampus at the apex of the visual hierarchy (Felleman and Van Essen, 1991). More recent non-human primate models (Kravitz et al., 2011) highlight multiple visual pathways originating in primary visual cortex that converge on the hippocampus, providing multiple routes for the feed-forward encoding of retinotopic information. Functional imaging studies have confirmed many features of this model in humans (Kravitz et al., 2011; Margulies et al., 2009; Silson et al., 2015) by demonstrating the contralateral encoding of visual field position in structures thought to be anatomically connected with the hippocampus (Margulies et al., 2009). Specifically, the scene-selective Parahippocampal Place Area (PPA), located in parahippocampal gyrus (PHG) (Epstein & Kanwisher, 1998), and Medial Place Area (MPA), located in medial parietal cortex (Silson et al., 2016) both exhibit biases for contralateral visual space.

Beyond the retinotopic nature of inputs to the hippocampus, a handful of studies provide neurophysiological support for retinotopic sensitivity in medial temporal lobe structures. For example, visually responsive cells have been recorded from the hippocampus and neighboring structures of non-human primates (Maclean et al., 1968; Desimone & Gross., 1979), and early electrophysiological recordings from the human hippocampal formation reported a pair of units with receptive fields in the contralateral upper visual field (Wilson et al., 1983). One recent fMRI study asked whether distinct regions of the hippocampus were associated with spatial memory relating to coarse-grained locations of the visual field (Jeye et al. 2018), however their focus was on memory - they did not perform retinotopic mapping or test for a main effect of visual field location, and a nuanced pattern of results was found.

Given the evidence for retinotopically organized input, we predicted that human hippocampus would exhibit a contralateral bias during population receptive field (pRF) mapping. We tested this prediction directly, by estimating pRFs using fMRI (Dumoulin & Wandell, 2008) in a sample of individual participants (n=27). Consistent with our predictions, a significant contralateral bias was present in bilateral hippocampus at the group-level. Further, the distribution of retinotopically sensitive voxels within the hippocampus highlighted a more prominent representation in anterior and medial portions. Finally, this contralateral bias was confirmed in an independent 7.0 Tesla retinotopy data set, collected as part of Human Connectome Project initiative (Benson et al., 2018; Szinte & Knapen, 2020).

## Materials and Methods

### Participants

Twenty-nine participants completed the initial fMRI experiment (21 females, mean age = 24.2 years). All participants had normal or corrected to normal vision and gave written informed consent. The National Institutes of Health Institutional Review Board approved the consent and protocol. This work was supported by the Intramural Research program of the National Institutes of Health – National Institute of Mental Health Clinical Study Protocols 93-M-0170, NCT00001360. (A further 181 participants were included in the HCP data set, detailed below.)

### fMRI scanning parameters

Participants were scanned on a 3.0T GE Sigma MRI scanner using a 32-channel head coil in the Clinical Research Center on the National Institutes of Health campus (Bethesda, MD). Across all participants, whole brain coverage was acquired. Slices were orientated axially, such that the most inferior slice was below the temporal lobe. All participants completed six population receptive field mapping runs and six runs of a six category-localizer. All functional images were acquired using a BOLD-contrast sensitive standard EPI sequence (TE = 30 ms, TR = 2 s, flip-angle = 65 degrees, FOV = 192 mm, acquisition matrix = 64×64, resolution 3 × 3 × 3 mm, slice gap = 0.3 mm, 28 slices). A high-resolution T1 structural image was obtained for each participant (TE = 3.47 ms, repetition time = 2.53 s, TI = 900 ms, flip angle = 7°, 172 slices with 1 x 1 x 1 mm voxels).

### Visual Stimuli and Tasks

#### Population receptive field mapping

During pRF mapping sessions a bar aperture traversed gradually through the visual field, whilst revealing randomly selected scene fragments from 90 possible scenes. During each 36 s sweep, the aperture took 18 evenly spaced steps every 2 s (1 TR) to traverse the entire screen. Across the 18 aperture positions all 90 possible scene images were displayed once. A total of eight sweeps were made during each run (four orientations, two directions). Specifically, the bar aperture progressed in the following order for all runs: Left to Right, Bottom Right to Top Left, Top to Bottom, Bottom Left to Top Right, Right to Left, Top Left to Bottom Right, Bottom to Top, and Top Right to Bottom Left. The bar stimuli covered a circular aperture (diameter = 20° of visual angle). Participants performed a color detection task at fixation, indicating via button press when the white fixation dot changed to red. Color fixation changes occurred semi-randomly, with approximately two-color changes per sweep (Silson et al., 2015). Stimuli for this and the other in-scanner task were presented using PsychoPy software (<u>Peirce, 2007</u>) (RRID:<u>SCR 006571</u>) from a Macbook Pro laptop (Apple Systems, Cupertino, CA).

### Six category functional localizer

Participants completed six functional localizer runs. During each run, color images from six stimulus categories (Scenes, Faces, Bodies, Buildings, Objects and Scrambled Objects) were presented at fixation (5 × 5° of visual angle) in 16 s blocks (20 images per block [300 ms per image, 500 ms blank]). Each category was presented twice per run, with the order of presentation counterbalanced across participants and runs. Participants responded via a MRI compatible button box whenever the same image appeared sequentially.

### fMRI data processing

#### Preprocessing

All data were analyzed using the Analysis of Functional NeuroImages (AFNI) software package (Cox, 1996) (RRID:SCR_005927). All functions and programs are readily available in the current version: AFNI binary version April 21, 2020. Before pRF and functional localizer analyses, all images for each participant were motion corrected to the first image of the first run (*3dVolreg*), after removal of the appropriate “dummy” volumes (eight) to allow stabilization of the magnetic field. Post motion-correction data were detrended (*3dDetrend*) and, in the case of the localizer data, smoothed with a 5 mm full-width at half-maximum Gaussian kernel (*3dMerge*).

### Population receptive field modelling

Detailed description of the pRF model implemented in AFNI is provided elsewhere (Silson et al., 2015). Briefly, given the position of the stimulus in the visual field at every time point, the model estimates the pRF parameters that yield the best fit to the data: pRF center location (x, y), and size (diameter of the pRF). Both Simplex and Powell optimization algorithms are used simultaneously to find the best time-series/parameter sets (x, y, size) by minimizing the least-squares error of the predicted time-series with the acquired time-series for each voxel.

### Six category functional localizer

Analyses were conducted using a general linear model approach and the AFNI programs *3dDeconvolve* and *3dREMLfit*. The data at each time point were treated as the sum of all effects thought to be present at that time point and the time series was compared against a Generalized Least Square (GLSQ) model fit with REML estimation of the temporal autocorrelation structure. Specifically, a response model was built by convolving a standard gamma function with a 16 s square wave for each condition and compared against the activation time courses using GLSQ regression. Motion parameters and four polynomials accounting for slow drifts were included as regressors of no interest. To derive the response magnitude per condition, *t*-tests were performed between the condition-specific beta estimates (normalized by the grand mean of each voxel for each run) and baseline.

### Anatomical Alignment

In each participant, both the pRF and functional localizer data were first de-obliqued (*3dWarp*) before being aligned to the individual participant’s high-resolution T1-weighted anatomical scan (*align_epi_anat.py*). Each participant’s aligned data were then inspected visually to confirm alignment accuracy. Given prior work demonstrating that the collateral sulcus (Weiner et al., 2018) and the mid-fusiform sulcus (Weiner et al., 2014) provide accurate anatomical landmarks for the peak of scene-selective PPA and face-selective Fusiform Face Area (FFA; Kanwisher et al., 1997), the results of the contrast Scenes versus Faces were overlaid onto each individual participants’ anatomical scan and inspected. Accurate alignment was determined using the above criteria for 27/29 participants. Subsequent analyses included only the 27 participants who met this alignment criteria.

### Hippocampal definitions

For each participant, the automated hippocampal segmentation provided by the output of Freesurfer 4 autorecon script (http://surfer.nmr.mgh.harvard.edu/) was used as a mask for the hippocampus. In order to divide the hippocampus into anterior, middle and posterior sections we first sorted the voxel indices by the y-axis, which codes for cortical anterior-posterior position. These indices were then separated into equal thirds and the corresponding pRF parameters were sampled for further analysis.

### Visual field coverage and visual field biases

The visual field coverage plots represent the group average sensitivity of each region of interest (ROI) to different positions in the visual field. To compute these, individual participant visual field coverage plots were first derived. These plots combine the best Gaussian receptive field model for each voxel within an ROI. Here, a max operator was used that reflects, at each point in the visual field, the maximum value from all pRFs within the ROI (Winawer et al., 2010). To compute visual field biases in individual participants and ROIs, we calculated the mean pRF sensitivity in the Ipsilateral and Contralateral visual fields, respectively.

### Statistical analyses

Statistics were calculated using the R Studio package (version 1.3). For our analyses, we used repeated-measures ANOVAs to examine the presence of contralateral biases in the hippocampus. For each analysis, we established initially whether the ANOVA adhered to the assumptions of sphericity using Mauchly’s test. When the assumption of sphericity was violated, the degrees of freedom for that main effect or interaction were corrected using the Greenhouse– Geisser correction to allow appropriate interpretation of the *F* value resulting from the ANOVA.

### HCP Retinotopy data

To confirm the contralateral biases in the hippocampus, we turned to the 7.0 Tesla retinotopy data set collected as part of the HCP initiative (Benson et al., 2018). This data set comprises high-resolution retinotopic data (1.6mm isotropic) and a large sample size (n=181). Full descriptions of this data set are provided elsewhere (Benson et al., 2018), but briefly, participants completed six retinotopic mapping runs (2x rotating wedge, 2x expanding ring 2x moving bar) in which the stimulus aperture presented a dynamic color texture (comprised of objects at different scales) on a pink noise background. Participants fixated centrally and indicated via button press when the fixation dot changed color. For consistency with our individual participant analyses we sampled the averaged data for the two bar runs only. Specifically, we sampled pRFs in the hippocampus from the group-averaged data derived by first computing the average time-course for each voxel across participants and, second, fitting the linear Gaussian pRF model to these group-averaged time-courses using custom python-based routines. Note that the pRF modelling implementation applied to the HCP data is different from that applied to the single participant data. Preprocessing on these data was identical to that used for the previous demonstration of retinotopic sensitivity within the default mode network (Szinte & Knapen, 2020). A mask for the hippocampus was taken from the Harvard/Oxford probabilistic atlas (https://fsl.fmrib.ox.ac.uk/fsl/fslwiki/Atlases).

## Results

We tested the hypothesis that the human hippocampus would exhibit a spatial bias for the contralateral visual field during visual field mapping **(Figure 1A)**. Such a bias would mirror not only early visual cortex, but also, more anterior regions, such as medial parietal cortex and parahippocampal cortex that provide input to the hippocampus both directly and indirectly.

**Figure 1.**
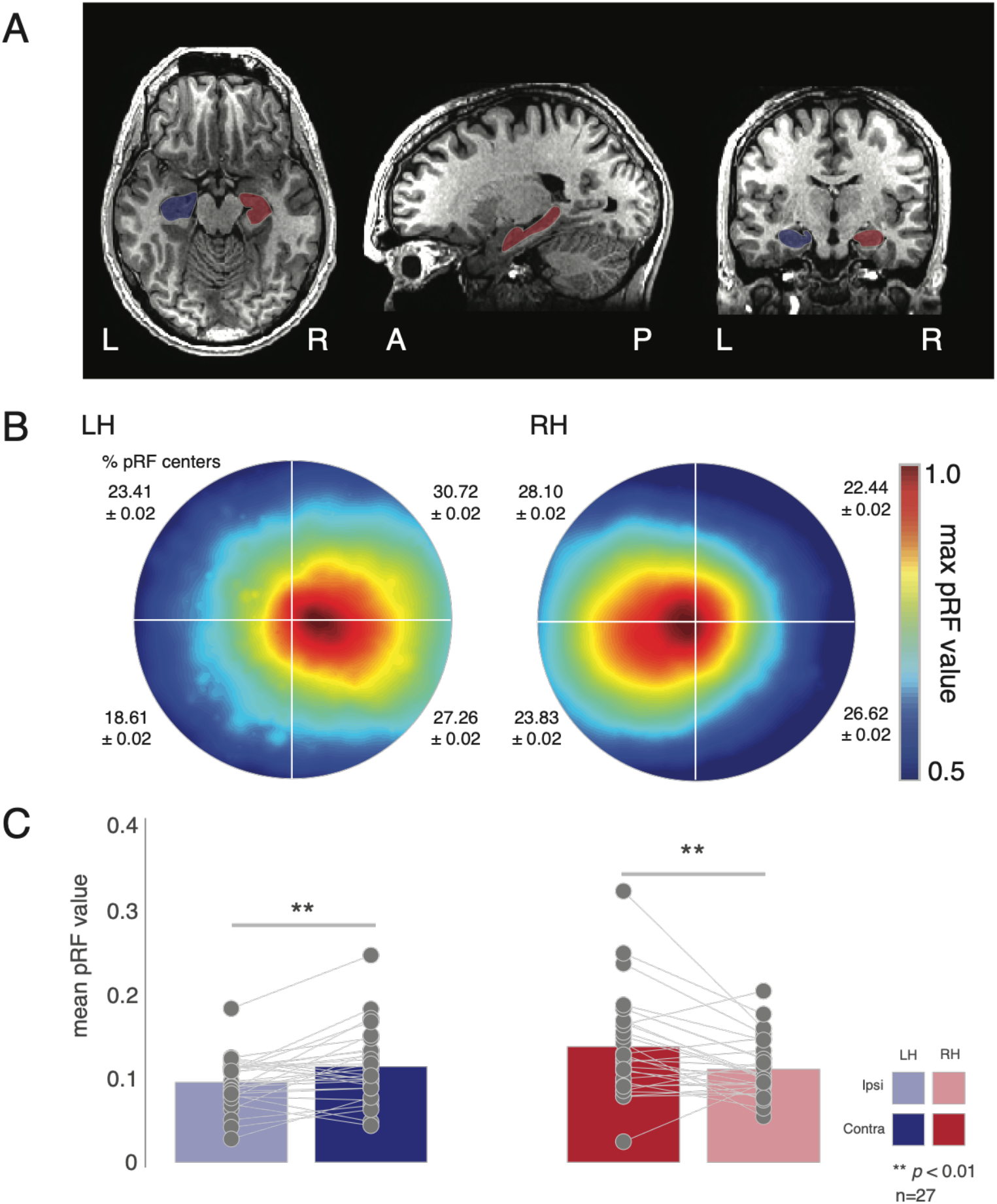
Contralateral biases in human hippocampus. **A,** Masks of the left (blue) and right (red) hippocampus of a representative participant. Images are in neurological convention. **B,** Group average (n=27) visual field coverage plots derived from all suprathreshold (R^2^ >= 0.1) voxels. A clear contralateral bias is evident in bilateral hippocampus. The mean percentage and standard deviation of pRF centers in each quadrant is shown inset. **C,** Quantification of contralateral biases. Bars represent the group-average pRF value in the ipsilateral (faded bars) and contralateral (solid bars) visual fields. Individual participant values are plotted and linked for each hippocampus. On average a significant contralateral bias was present in both hemispheres. **p<0.01.

### Biased representation of contralateral space in the hippocampus

Initially, we computed visual field coverage plots in each participant and ROI (left hippocampus, right hippocampus) from all suprathreshold pRFs (R^2^ >= 0.1), before averaging these coverage plots across participants. These visual field coverage plots represent schematic visualizations of the sensitivity of a given brain region to different positions in the visual field, built by combining the best Gaussian receptive field model (position, size and explained variance) for each voxel within an ROI. In our analyses, a max operator is used. This creates a coverage plot that reflects, at each point in the visual field, the maximum sensitivity (which we refer to as pRF value) from all of the receptive field models within an ROI (min=0, max=1) Thus, the coverage plot reflects the maximum envelope of all the pRFs.

The group average visual field coverage plots for the left and right hippocampus **(Figure 1B)** demonstrate a striking contralateral bias for both hemispheres, respectively. From the average coverage plots alone, there is no clear evidence of any quadrant biases but note the numerically higher percentages of pRF centers in the upper visual field **(inset Figure 1B)**.

To quantify these contralateral biases, we calculated the mean pRF value (see above) in the ipsilateral and contralateral visual field in each participant and ROI, respectively, and submitted these to a two-way repeated measures ANOVA with Hemisphere (Left, Right) and Visual Field (Ipsilateral, Contralateral) as within-participant factors. The main effects of Hemisphere (F_(1, 26)_ = 6.98, *p*=0.02, partial eta^2^=0.06) and Visual field (F_(1, 26)_ = 21.44, *p*=8.89^−5^, partial eta^2^=0.07) were significant, reflecting on average larger pRF values in the right over left hemisphere and the contralateral over ipsilateral visual field, respectively. The Hemisphere by Visual Field interaction was not significant (*p*>0.05). A series of paired *t*-tests confirmed a significant contralateral bias in both the left (*t*_(26)_=2.50, *p*=0.01) and right (t_(26)_=3.22, *p*=0.003) hippocampus **(Figure 1C)**.

### Retinotopic sensitivity and scene-selectivity in the hippocampus

Given prior work suggesting a place for the hippocampus in the scene-processing network (Maguire & Mullally, 2013; Hodgetts et al., 2016), we next sought to establish the relationship between the strength of retinotopic encoding (variance explained by the pRF model) and the degree of scene-selectivity within the hippocampus. In each participant and hemisphere, we calculated the correlation (Pearson’s) between the variance explained by each voxel’s pRF fit and that voxel’s corresponding index of scene-selectivity (*t-value* of the contrast Scenes versus Faces in a separate localizer task), before averaging correlation coefficients across participants. On average, a positive correlation was observed in each hemisphere, suggesting that the more retinotopically sensitive a voxel, the more scene-selective also **(Figure 2A)**. A series of *t*-tests versus zero (i.e. no correlation) confirmed the significant positive correlation at the group level in both hemispheres (lh: t_(26)_=3.88, *p*=0.002, rh: t_(26)_=3.23, *p*=0.003).

**Figure 2.**
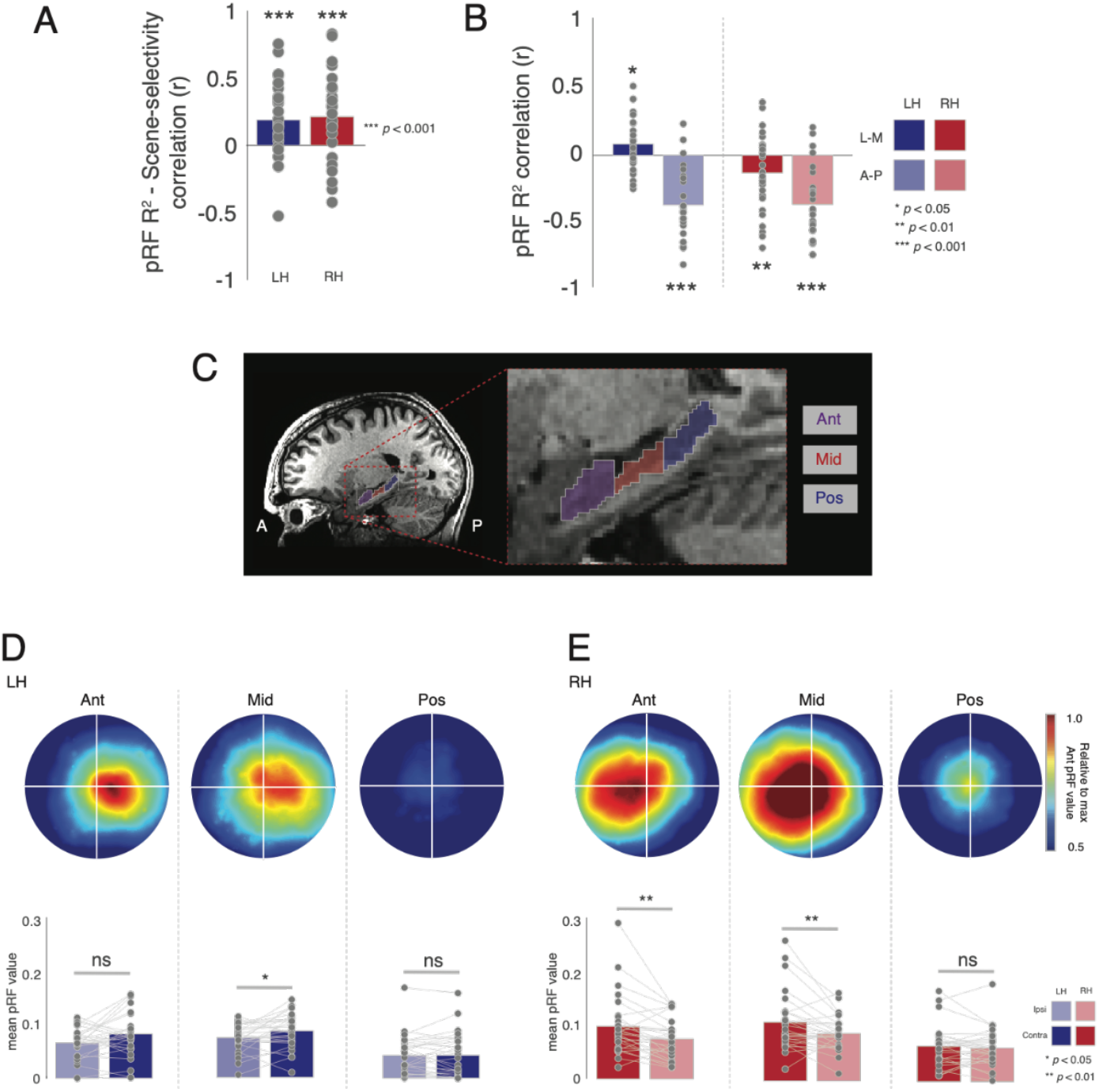
Relationship with scene-selectivity and contralateral biases in hippocampal sections. **A,** Bars represent the group-average correlation (Pearson’s) between pRF R^2^ and scene-selectivity across voxels. **B,** Bars represent the group average correlation between pRF R^2^ and position along the lateral-medial (solid bars) and anterior-posterior (faded bars) axes. **C,** Enlarged view of the hippocampus showing the Anterior, Middle and Posterior sections. **D,** Group average visual field coverage plots derived from all suprathreshold (R^2^ > 0.1) voxels in each hippocampal section. **D,** Bars represent the group-average pRF value in the ipsilateral (faded bars) and contralateral (solid bars) visual fields for each section in the left hippocampus. **E**, same as D but for the right hippocampus. *p<0.05, **p<0.01.

### Distribution of retinotopic sensitivity within the hippocampus

Prior work suggests functional differences throughout the hippocampus, and a particularly common finding has been scene-selective responses in the medial (rather than lateral) aspect of anterior hippocampus (Zeidman & Maguire, 2016). To explore the spatial distribution of retinotopic sensitivity within the hippocampus, we sorted the voxel indices of each hippocampus first by the x-axis, which codes for left-right, and then by the y-axis, which codes for anterior-posterior within the brain. Next, we computed the correlation (Pearson’s) between each voxel’s position along that axis and the strength of retinotopic encoding (pRF explained variance), before averaging correlation coefficients across participants and testing against zero (i.e. no correlation) **(Figure 2B)**. In both hemispheres, there was a significant correlation between absolute x-position and retinotopic sensitivity (lh: t_(26)_=2,41, *p*=0.02, rh: t_(26)_=2.26, *p*=0.03), reflecting better pRF model fits medially, as well as, significant negative correlations between y-position and retinotopic sensitivity, reflecting greater explained variance anteriorly (lh: t_(26)_=7.52, *p*=5.42^−8^, rh: t_(26)_=7.91, 2.17^−8^).

We next sought to establish whether a contralateral bias would be present in sub-sections of the hippocampus. Accordingly, we divided each participant’s hippocampus into equal thirds along the y-axis (see methods). These were subsequently labelled as Anterior, Middle and Posterior sections **(Figure 2C)**. The group average visual field coverage plots for each section are depicted for the left **(Figure 2D)** and right hippocampus **(Figure 2E)**. At the group level, a clear contralateral bias is evident in the anterior and middle sections of both hemispheres, whereas the posterior sections exhibit no such bias.

To quantify these biases, we computed the mean pRF value in both the ipsilateral and contralateral visual fields in each individual participant and ROI. These values were submitted to a three-way repeated measures ANOVA with Hemisphere (Left, Right), Section (Anterior, Middle, Posterior) and Visual Field (Ipsilateral, Contralateral) as within-participant factors. The main effects of Hemisphere (F_(1, 26)_=8.75, *p*=0.006, partial eta^2^=0.05), Section (F_(2, 52)_=23.49, *p*=5.38^−8^, partial eta^2^=0.16) and Visual Field (F^(1, 26)^=20.14, *p*=0.0001, partial eta^2^=0.02), were significant, reflecting on average larger pRF values in the right hemisphere, in anterior and middle over posterior sections and in the contralateral over ipsilateral visual field, respectively. Only the Section by Visual field interaction (F_(2, 52)_=5.75, *p=*0.01, partial eta^2^=0.008, GG-corrected) was significant. All other interactions were not significant (*p*>0.05, in all cases).

To explore this further, we conducted a series of two-way ANOVAs with Section and Visual Field as factors in each hemisphere separately. In the left hemisphere, only the main effect of Section (F_(2, 52)_=25.58, *p*=1.83-8, partial eta^2^=0.21) was significant (*p*>0.05, in all other cases). A series of paired *t*-tests revealed a significant contralateral bias in the middle (t_(26)_=1.96, *p*=0.02), but not the anterior (t_(26)_=1.61, *p*=0.10) or posterior sections (t_(26)_=0.14, *p*=0.44), although note the numerically larger contralateral bias in the anterior section **(Figure 2F)**. In the right hemisphere, both the main effects of Section F_(2, 52)_=11.28, *p*=8.51^−5^, partial eta^2^=0.12) and Visual field (F_(2, 52)_=9.99, *p*=0.003, partial eta^2^=0.04) were significant, as was their interaction (F_(2, 52)_=5.52, *p*=0.01, partial eta^2^=0.01, GG-corrected). Again, a series of paired *t-*tests revealed significant contralateral biases in both the anterior (t_(26)_=4.00, *p*=0.0004) and middle (t_(26)_=2.88, *p*=0.007), but not the posterior section (t_(26)_=0.96, *p*=0.34) **(Figure 2G)**.

### Reduced signal posteriorly could explain lack of contralateral bias

Whilst the data suggest that the strength of retinotopic sensitivity is reduced more posteriorly in the hippocampus, it is important to consider the impact of signal strength on these patterns of results. First, we calculated the temporal signal-to-noise (tSNR) of the pRF runs for each participant. Next, we computed the median tSNR values in each section of the hippocampus and submitted these values to a two-way repeated measures ANOVA with Hemisphere and Section as factors (same levels as above). The main effect of Section was significant (F_(2, 52)_=110.54, *p*=1.54^−13^, partial eta^2^=0.40, GG-corrected), reflecting larger tSNR values more anteriorly, whereas the main effect of Hemisphere and the Hemisphere by Section interaction were not significant (*p*>0.05 in both cases). Given the non-significant effect of Hemisphere, tSNR values were averaged across hemispheres before being submitted to a one-way ANOVA with Section as the only factor. The main effect of Section was significant (F_(2, 52)_=110.54, *p*=1.54^−13^, partial eta^2^=0.43, GG-corrected). A series of paired *t-*tests confirmed that tSNR decreased significantly from anterior to posterior in the hippocampus (Anterior *versus* Middle: t_(26)_=10.56, *p*=6.65^−11^; Anterior *versus* Posterior: t_(26)_=11.32, *p*=1.49^−11^; Middle *versus* posterior: t_(26)_=8.16, *p*=1.19^−8^).

### Contralateral bias in hippocampus not due to spillover from PHG

The hippocampus is located anterior and dorsal of the parahippocampal gyrus (PHG). Prior work from our group and others has demonstrated the strong influence of retinotopy in the parahippocampal gyrus and in the PPA in particular. Given the known proximity between the PHG and the hippocampus we sought to rule out the possibility that these retinotopically sensitive responses measured within the hippocampus were due to spillover of responses from PHG. In each participant, we examined the responses within the hippocampus with respect to those measured from PHG. The explained variance of the pRF model for a representative participant is shown in **Figure 3 (top)**. Whilst robust fits to the pRF model are evident in early visual cortex, extending anteriorly into ventral temporal cortex and encompassing the PPA, two small clusters of suprathreshold voxels are also evident within the hippocampus. These clusters, particularly the more anterior cluster, are spatially separated from responses in ventral temporal cortex and are unlikely to reflect spillover from PHG. Both clusters exhibit pRF centers located well within the contralateral visual field **Figure 3 (bottom)**.

**Figure 3.**
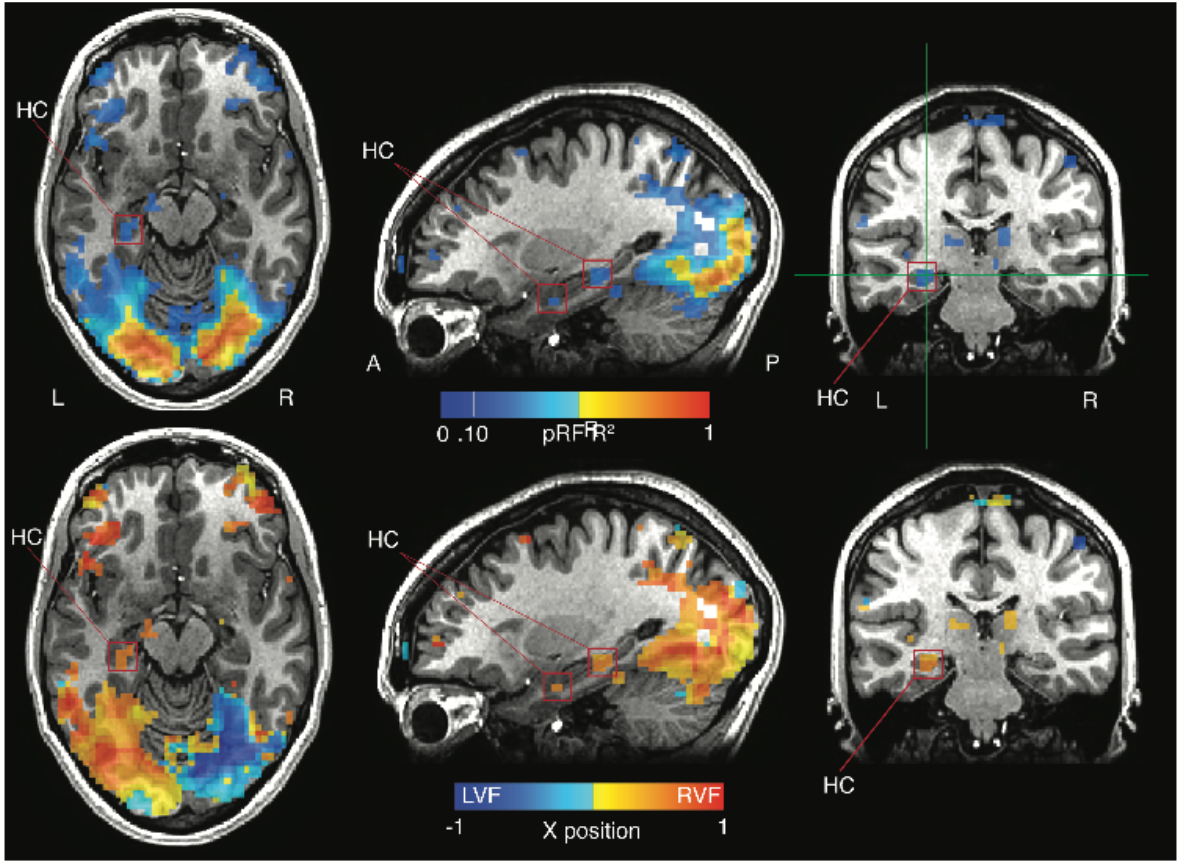
Retinotopic sensitivity in the hippocampus is spatially separate from PHG. **Top row,** The pRF R^2^ is overlaid onto axial, saggital and coronal slices of a representative participant. Strong responses are evident throughout visual cortex and extend anteriorly in ventral temporal cortex. Two clusters within the hippocampus (red boxes) appear spatially distinct from more posterior responses in PHG. **Bottom row,** The x-position of pRF centers are overlaid onto the same slices. The two hippocampal clusters show pRF positions firmly in the contralateral (right) visual field.

### Replication of contralateral bias in a high-resolution independent dataset

Our individual participant analyses demonstrate that, when considered as a single structure, the human hippocampus exhibits a significant bias for contralateral visual space when measured through pRF mapping. We next sought to confirm these findings in independent data by taking advantage of the large sample (n=181) and high-resolution (1.6mm isotropic) 7.0 Tesla retinotopy data collected as part of the HCP initiative (Benson et al., 2018).

Using the group average pRF fitted data (from the bar runs only), we sampled pRF parameters (R^2^, x-position, eccentricity and pRF size) from a mask of the hippocampus. Enlarged views of the hippocampus with each pRF parameter overlaid in false colour are shown in **Figure 4**. Many of the features present in the individual participant data are also evident here, despite these data being acquired across different scanners, fieldstrengths, resolutions and visual stimulus setups, while also being analysed using different processing pipelines. These data demonstrate **a)** that voxels are fit well by the pRF model throughout the hippocampus, with clear clusters evident in anterior medial sections **(Figure 4A)**, **b)** hippocampal pRFs exhibit largely contralateral visual field centers **(Figure 4B)**, **c)** pRFs are relatively eccentric with few representing the fovea and **d)** pRFs range is size but with very few small pRFs. For completeness, we calculated the visual field coverage in each hemisphere from all supratheshold pRFs (R^2^ > 0.1) from the HCP data. In both hemispheres, a clear contralateral bias is evident **(Figure 4E)**. Again, there is no clear evidence for any quadrant biases but note that unlike our individual participant analyses the HCP data contains a higher percentage of lower visual field centers (percentage of pRF centers inset). These data complement the individual participant analyses reported above and highlight the contralateral bias exhibited by the human hippocampus during visual field mapping.

**Figure 4.**
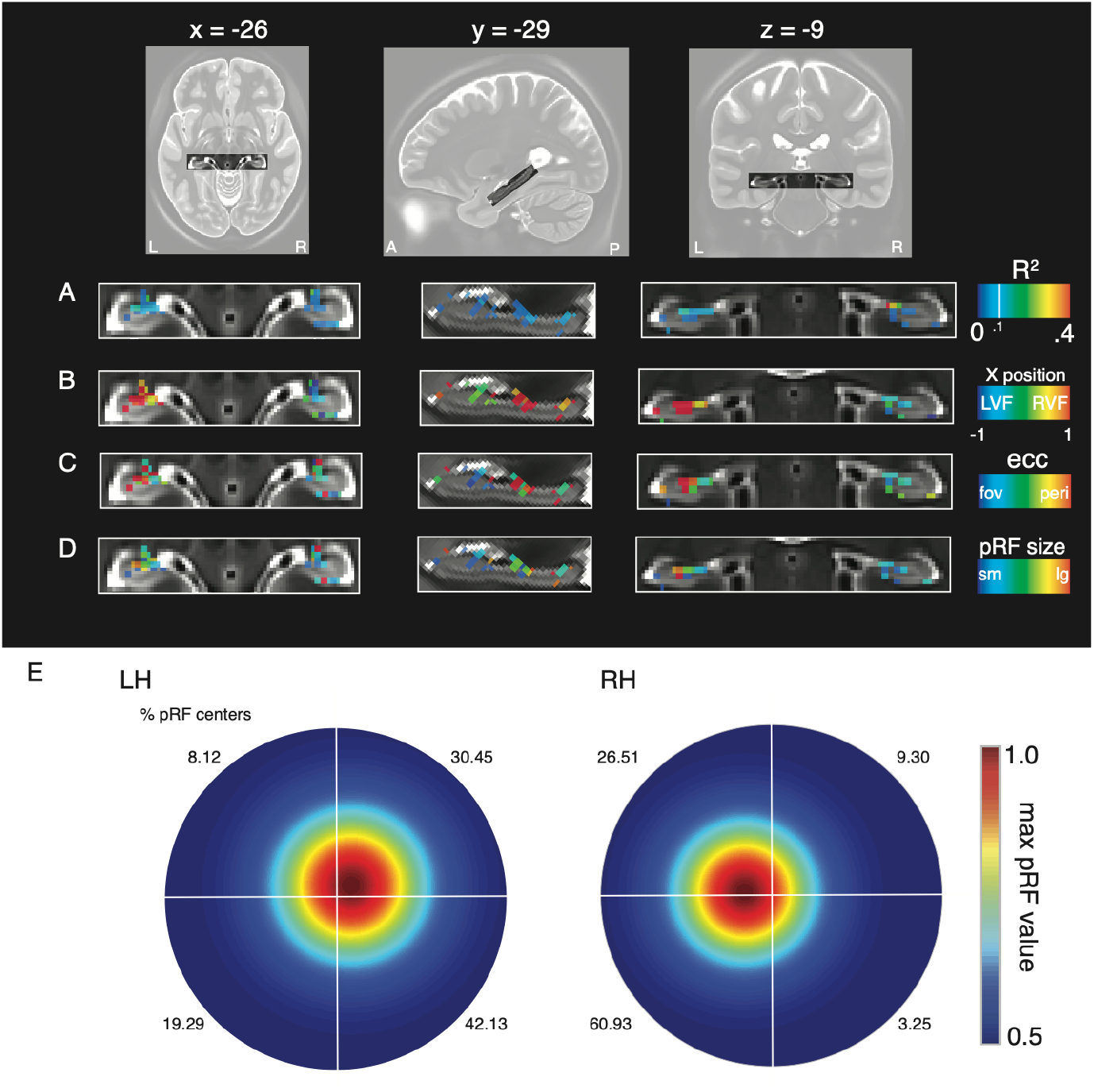
pRF parameters in the hippocampus from the HCP data. **A,** Enlarged axial, sagittal and coronal views of the hippocampus shown with the pRF R^2^ overlaid. Voxels in the hippocampus are well fitted by the pRF model, with clusters in anterior medial portions. **B,** The x-position of pRFs is shown. In general, pRFs show largely contralateral visual field positions. **C,** pRF eccentricity suggests peripheral pRFs in the hippocampus. **D,** Hippocampal pRFs appear also to be large. **E,** Visual field coverage from all suprathreshold pRFs (R^2^ > 0.1; left = 199, right = 115). A contralateral bias is present in bilateral hippocampus. The percentage of pRF centers in each quadrant are inset.

## Discussion

Here, using pRF data from two independent sources we demonstrate a consistent contralateral bias in the human hippocampus during visual field mapping. These data demonstrate that the influence of retinotopy is present and measurable even at the very highest level of the visual hierarchy (Fellemen and Van Essen, 1991; Kravitz et al., 2011; 2013) and suggests that retinotopy be considered as a visuospatial representation that is available to the hippocampus.

### Anatomical connectivity with the hippocampus implies retinotopic sensitivity

A contralateral bias of visual space was implied by direct and indirect connections between the hippocampus and antecedent regions of the visual hierarchy. Tract-tracing studies in non-human primates positioned the hippocampus at the highest-level in the visual hierarchy (Fellemen and Van Essen, 1991). The regions with which it is connected are responsive to visual stimuli, with early neurophysiological studies identifying visually responsive units in parahippocampal structures in non-human primates (Maclean et al., 1968; Desimone & Gross., 1979) and humans (Wilson et al., 1983). More recently, functional neuroimaging has demonstrated contralateral population receptive fields in multiple regions thought to connect directly and/or indirectly with the hippocampus (Silson et al., 2015). Specifically, contralateral biases have been reported in scene-selective PPA, OPA and MPA – located on the ventral, lateral and medial surfaces, respectively (Silson et al., 2015; 2016).

We found that retinotopic population receptive fields were detectable in the human hippocampus, lateralized to each contralateral hemisphere, using two independent datasets with distinct stimuli. We did not find any evidence for a systematic mapping of visual space in the hippocampus - a hallmark of early visual cortex. However, the absence of a retinotopic map should not imply the absence of retinotopic sensitivity. Indeed, prior work from our group (Silson et al., 2015; 2016) and others (Elshout et al., 2018) has demonstrated robust and reliable retinotopically driven responses in occipitotemporal and medial parietal cortices without clear evidence for accompanying retinotopic maps. Moreover, this could be due to technical limitations: given the organizational scale of the hippocampus relative to current fMRI voxel sizes it is possible that finding map-like organization in hippocampus requires using even smaller fMRI voxels. The coarse representation of contralateral visual space reported here is consistent with a very recent study employing ultra-high resolution and connective field modelling to demonstrate fine-grained visuotopic connectivity between V1 and the hippocampus (Knapen, 2020). The question of whether the contralateral biases reported here (and elsewhere, Knapen, 2020) reflect retinotopic inputs into the hippocampus or retinotopic neurons within the hippocampus itself cannot be answered by the current fMRI data, but remains an important and open question for future research.

### Distribution of retinotopic sensitivity across the hippocampus

Studies of visual scene perception and discrimination have highlighted the potentially key role played by the anterior medial portion of the hippocampus (Hogetts et al., 2016; Zeidman and Maguire, 2016). Our results were consistent with this. Not only did we observe, on average, a significant positive correlation between retinotopic sensitivity and medial – lateral position within the hippocampus, but also, a significant negative correlation between retinotopic sensitivity and anterior-posterior position. Subsequent analyses of separate hippocampal sections also suggested more prominent retinotopic sensitivity anteriorly, but these are to be interpreted with caution as follow-up analyses also revealed that tSNR drops systematically in more posterior regions.

Our data demonstrated a significant positive relationship between retinotopic sensitivity and scene-selectivity, suggesting that the well-established preferential response of the hippocampus to scene stimuli involves processing in retinotopic space. Interestingly, similar positive relationships between scene-selectivity and retinotopy have been reported within scene-selective MPA in medial parietal cortex (Silson et al., 2016), which is thought to provide input to the hippocampus (Margulies et al., 2009; Kravitz et al., 2011).

### Visuospatial encoding in the hippocampus

What information might the hippocampus be encoding or processing? The hippocampus directly encodes an animal’s spatial location in an allocentric (world-centered) reference frame (O’Keefe, and Dostrovsky, 1971). Visual input contributes to the formation of these representations (Chen et al., 2013), and indeed, recent findings have demonstrated that neuronal populations in both CA1 and V1 encode the rodent’s subjective estimate of its position along a linear track (Saleem et al., 2018). However, to our knowledge, retinotopy has never been identified in the rodent hippocampus, which may be unsurprising given their large, overlapping visual fields and relatively poor visual acuity.

There is increasing evidence that primate hippocampus and entorhinal cortex encode not only physical location, but also *visual space* in multiple reference frames (Miester, 2018; Rolls and Wirth, 2018; Zeidman and Maguire, 2016; Nau et al., 2018). In brief, primate *spatial view cells* were found to encode positions on a video screen, or the position of the video screen in the room (Feigenbaum and Rolls, 1991; Georges-François, Rolls & Robertson, 1999). More recently, entorhinal grid cells (Hafting et al., 2005) have been found to have firing fields covering gaze direction or visual space in non-human primates (Killian et al., 2012; Wilming et al., 2018) and in humans (Nau et al., 2018; Julian et al., 2018). Our results demonstrate that retinotopy complements these other visuospatial representations in the hippocampus.

### Functional significance of multiple visuospatial representations

What functions might be served by the presence of multiple visuospatial representations in the hippocampus? Insights may be gained from neuropsychological studies on patients with specific lesions to the hippocampus. Such patients have been found to be impaired at discriminating images of similar three-dimensional scenes, or scenes from different viewpoints (Lee et al., 2005; 2005; Aly et al., 2013; Suzuki et al., 2009; Baxter et al., 2009) and they are impaired at extrapolating beyond the view (Mullally et al., 2012). Thus, the hippocampus may be required for complex visual tasks, which require forming an internal representation or model of the stimuli. Our results suggest this may be subserved by conjunctive retinotopic and allocentric representations in the hippocampus.

Neuropsychological theories have been proposed to explain these findings in patients. In particular, *scene construction theory* (Hassabis et al., 2007) proposes that the hippocampus and connected regions form internal models of scenes, facilitating cognitive functions including vision, navigation, imagination and episodic memory (Ziedmann & Maguire, 2013). Under this account, the hippocampus could be considered a node in the scene-processing network (Maguire & Mullally, 2013; Hodgetts et al., 2016), that is functionally connected to antecedent scene-selective regions (Margulies et al., 2009; Silson et al., 2016) and these regions exhibit prominent biases for contralateral visual space (Silson et al., 2015). Thus, the left hippocampus may contribute information from the right visual field to the formation of a scene representation, and vice versa. Our initial pRF modelling employed scene stimuli whereby multiple scene fragments were presented at each location. Whilst this paradigm was used to try and prevent participants from mentally ‘filling-in’ the scenes, it is possible that scene fragments were namable and generated internal representations. On the other hand, the stimulus employed under the HCP initiative (Benson et al., 2018) could be considered far more abstract (objects at multiple scales on a pink-noise background).

An alternative perspective on hemifield-specific responses recognizes that the hippocampus guides behaviour, and this behaviour may include eye movements. The level of hippocampus activity has been found to correlate with the number of fixations when novel face images are presented, suggesting a role for the hippocampus in sampling information (Liu et al., 2017). A recent proposal, the *spatiotemporal similarity hypothesis*, explains this by suggesting that that the hippocampus represents stimuli that co-occur in space and time, and it uses these joint representations to generate visual predictions and guide eye movements (Turk-Browne, 2019). *Predictive coding* is a computational framework which formalizes these notions and comes in multiple forms. Particularly relevant is *active inference* (Friston et al., 2015), which treats the brain as a deep hierarchical forward model that predicts sensory information and infers the causes of sensations by taking actions (such as sampling new information). Under this account, the purpose of a visual saccade is to test a hypothesis (i.e., reduce uncertainty) about what might be ‘out there’ beyond the current view (Parr and Friston, 2018). The contribution of the hippocampus is proposed to be encoding transitions between discrete states, such as sequences of eye gaze positions (Mirza et al., 2016). Our results might suggest that left hippocampus encodes potential sequences of eye movements related to the right visual field, and vice versa (although in the tasks we present here, any such motor plans could not be enacted, as subjects were required to fixate centrally).

Finally, the hippocampus may also encode temporal regularities, sequences or transition probabilities in the environment (Stachenfeld et al., 2017; Kumaran et al., 2006; Garvert et al., 2017). The pRF stimuli were highly predictable, traversing gradually on a predetermined trajectory through the visual field. It is therefore possible that the responses were elicited by predictions related to the sequence of stimuli in the contralateral visual field. An interesting future experiment could test this hypothesis by manipulating the predictability of the retinotopic mapping stimuli and measuring its impact on the contralateral biases measured as a result.

### Conclusion

Taken together, our data highlight that retinotopic sensitivity, and the contralateral encoding of visual information in particular, is present even at the level of the human hippocampus. Whether such sensitivity reflects retinotopic input or the activity of retinotopic neurons in the hippocampus remains unclear. Likewise, how the hippocampus incorporates this retinotopic information with the allocentric and global spatial representations that the hippocampus supports is an important goal of future work, but it is possible that such a representation provides a means for the hippocampus to compare ongoing sensory inputs with past events. Indeed, the seemingly ubiquitous encoding of retinotopic information within brain regions that subserve divergent functions suggests the brain may utilize retinotopy as a means to facilitate neural communication.

## Acknowledgments

We thank Matthias Nau for thoughtful comments on the manuscript.

## References

Aggleton, J.P. (2012). Multiple anatomical systems embedded within the primate medial temporal lobe: Implications for hippocampal function. Neurosci Biobehav R 36, 1579–1596.

Aly, M., Ranganath, C., and Yonelinas, A.P. (2013). Detecting Changes in Scenes: The Hippocampus Is Critical for Strength-Based Perception. Neuron 78, 1127–1137.

Battaglia, F.P., Sutherland, G.R., and McNaughton, B.L. (2004). Local sensory cues and place cell directionality: additional evidence of prospective coding in the hippocampus. Journal of Neuroscience 24, 4541–4550.

Baxter, M.G. (2009). Involvement of medial temporal lobe structures in memory and perception. Neuron 61, 667–677.

Bender, F., Gorbati, M., Cadavieco, M.C., Denisova, N., Gao, X., Holman, C., Korotkova, T., and Ponomarenko, A. (2015). Theta oscillations regulate the speed of locomotion via a hippocampus to lateral septum pathway. Nature communications 6, 1–11.

Benson, N. C., Jamison, K. W., Arcaro, M. J., Vu, A. T., Glasser, M. F., Coalson, T. S., & Kay, K. (2018). The Human Connectome Project 7 Tesla retinotopy dataset: Description and population receptive field analysis. Journal of vision, 18(13), 23–23.

Chan, A. W., Kravitz, D. J., Truong, S., Arizpe, J., & Baker, C. I. (2010). Cortical representations of bodies and faces are strongest in commonly experienced configurations. Nature neuroscience, 13(4), 417–418.

Chen, G., King, J.A., Burgess, N. and O’Keefe, J., 2013. How vision and movement combine in the hippocampal place code. Proceedings of the National Academy of Sciences, 110(1), pp.378–383.

Desimone, R., & Gross, C. G. (1979). Visual areas in the temporal cortex of the macaque. Brain research, 178(2-3), 363–380.

Ding, S.L. (2013). Comparative anatomy of the prosubiculum, subiculum, presubiculum, postsubiculum, and parasubiculum in human, monkey, and rodent. The Journal of comparative neurology 521, 4145–4162.

Dumoulin, S. O., & Wandell, B. A. (2008). Population receptive field estimates in human visual cortex. Neuroimage, 39(2), 647–660.

Elshout, J. A., van den Berg, A. V., & Haak, K. V. (2018). Human V2A: A map of the peripheral visual hemifield with functional connections to scene-selective cortex. Journal of vision, 18(9), 22–22.

Epstein, R., & Kanwisher, N. (1998). A cortical representation of the local visual environment. Nature, 392(6676), 598–601.

Felleman, D.J., and Van, D.E. (1991). Distributed hierarchical processing in the primate cerebral cortex. Cerebral cortex (New York, NY: 1991) 1, 1–47.

Feigenbaum, J.D., and Rolls, E.T. (1991). Allocentric and egocentric spatial information processing in the hippocampal formation of the behaving primate. Psychobiology 19, 21–40.

Friston, K., Rigoli, F., Ognibene, D., Mathys, C., Fitzgerald, T., and Pezzulo, G. (2015). Active inference and epistemic value. Cognitive neuroscience 6, 187–214.

Garvert, M.M., Dolan, R.J., and Behrens, T.E. (2017). A map of abstract relational knowledge in the human hippocampal–entorhinal cortex. Elife 6, e17086.

Georges-François, P., Rolls, E.T., and Robertson, R.G. (1999). Spatial view cells in the primate hippocampus: allocentric view not head direction or eye position or place. Cerebral cortex 9, 197–212.

Groen, I. I., Silson, E. H., & Baker, C. I. (2017). Contributions of low-and high-level properties to neural processing of visual scenes in the human brain. Philosophical Transactions of the Royal Society B: Biological Sciences, 372(1714), 20160102.

Hafting, T., Fyhn, M., Molden, S., Moser, M.-B., and Moser, E.I. (2005). Microstructure of a spatial map in the entorhinal cortex. Nature 436, 801–806.

Hassabis, D., and Maguire, E.A. (2007). Deconstructing episodic memory with construction. Trends in cognitive sciences 11, 299–306.

Hemond, C. C., Kanwisher, N. G., & De Beeck, H. P. O. (2007). A preference for contralateral stimuli in human object-and face-selective cortex. PLoS one, 2(6).

Hodgetts, C. J., Shine, J. P., Lawrence, A. D., Downing, P. E., & Graham, K. S. (2016). Evidencing a place for the hippocampus within the core scene processing network. Human brain mapping, 37(11), 3779–379.

Huang, R.-S., and Sereno, M.I. (2013). Bottom-up retinotopic organization supports top-down mental imagery. The open neuroimaging journal 7, 58.

Jeye, B. M., MacEvoy, S. P., Karanian, J. M., & Slotnick, S. D. (2018). Distinct regions of the hippocampus are associated with memory for different spatial locations. Brain research, 1687, 41–49.

Julian, J.B., Keinath, A.T., Frazzetta, G., and Epstein, R.A. (2018). Human entorhinal cortex represents visual space using a boundary-anchored grid. Nature neuroscience 21, 191–194.

Johnson, A., and Redish, A.D. (2007). Neural ensembles in CA3 transiently encode paths forward of the animal at a decision point. Journal of Neuroscience 27, 12176–12189.

Kanwisher, N., McDermott, J., & Chun, M. M. (1997). The fusiform face area: a module in human extrastriate cortex specialized for face perception. Journal of neuroscience, 17(11), 4302–4311.

Killian, N.J., Jutras, M.J., and Buffalo, E.A. (2012). A map of visual space in the primate entorhinal cortex. Nature 491, 761–764.

Kravitz, D. J., Kriegeskorte, N., & Baker, C. I. (2010). High-level visual object representations are constrained by position. Cerebral Cortex, 20(12), 2916–2925.

Kravitz, D. J., Saleem, K. S., Baker, C. I., & Mishkin, M. (2011). A new neural framework for visuospatial processing. Nature Reviews Neuroscience, 12(4), 217–230.

Kravitz, D. J., Saleem, K. S., Baker, C. I., Ungerleider, L. G., & Mishkin, M. (2013). The ventral visual pathway: an expanded neural framework for the processing of object quality. Trends in cognitive sciences, 17(1), 26–49.

Kumaran, D., and Maguire, E.A. (2006). An unexpected sequence of events: mismatch detection in the human hippocampus. PLoS biology 4.

Lee, A.C.H., Buckley, M.J., Pegman, S.J., Spiers, H., Scahill, V.L., Gaffan, D., Bussey, T.J., Davies, R.R., Kapur, N., Hodges, J.R., et al. (2005). Specialization in the medial temporal lobe for processing of objects and scenes. Hippocampus 15, 782–797.

Lee, A.C.H., Bussey, T.J., Murray, E.A., Saksida, L.M., Epstein, R.A., Kapur, N., Jr, H., and Graham, K.S. (2005). Perceptual deficits in amnesia: challenging the medial temporal lobe ‘mnemonic’ view. Neuropsychologia 43, 1–11.

Liu, Z.X., Shen, K., Olsen, R.K. and Ryan, J.D., 2017. Visual sampling predicts hippocampal activity. Journal of Neuroscience, 37(3), pp. 599–609.

MacLean, P. D., Yokota, Toshikatsu., & Kinnard, M. A. (1968). Photically sustained on-responses of units in posterior hippocampal gyrus of awake monkey. Journal of neurophysiology, 31(6), 870–883.

Mackey, W. E., Winawer, J., & Curtis, C. E. (2017). Visual field map clusters in human frontoparietal cortex. Elife, 6, e22974.

Maguire, E.A. and Mullally, S.L., 2013. The hippocampus: a manifesto for change. Journal of Experimental Psychology: General, 142(4), p.1180.

Malcolm, G. L., Groen, I. I., & Baker, C. I. (2016). Making sense of real-world scenes. Trends in Cognitive Sciences, 20(11), 843–856.

Meister, M. (2018). Memory system neurons represent gaze position and the visual world. Journal of experimental neuroscience 12, 1179069518787484.

Mirza, M.B., Adams, R.A., Mathys, C.D., and Friston, K.J. (2016). Scene construction, visual foraging, and active inference. Frontiers in computational neuroscience 10, 56.

Momennejad, I., Russek, E.M., Cheong, J.H., Botvinick, M.M., Daw, N.D., and Gershman, S.J. (2017). The successor representation in human reinforcement learning. Nature Human Behaviour 1, 680–692.

Mullally, S.L., Intraub, H., and Maguire, E.A. (2012). Attenuated Boundary Extension Produces a Paradoxical Memory Advantage in Amnesic Patients. Current Biology 22, 261–268.

Nau, M., Julian, J. B., & Doeller, C. F. (2018). How the brain’s navigation system shapes our visual experience. Trends in cognitive sciences, 22(9), 810–825.

Nau, M., Schröder, T.N., Bellmund, J.L., and Doeller, C.F. (2018). Hexadirectional coding of visual space in human entorhinal cortex. Nature neuroscience 21, 188–190.

O’Keefe, J., and Dostrovsky, J. (1971). The hippocampus as a spatial map. Preliminary evidence from unit activity in the freely-moving rat. Brain research 34, 171–175.

Parr, T., and Friston, K.J. (2018). The discrete and continuous brain: from decisions to movement—and back again. Neural Comput 30, 2319–2347.

Poppenk, J., Evensmoen, H. R., Moscovitch, M., & Nadel, L. (2013). Long-axis specialization of the human hippocampus. Trends in cognitive sciences, 17(5), 230–240.

Race, E., Keane, M.M., and Verfaellie, M. (2011). Medial Temporal Lobe Damage Causes Deficits in Episodic Memory and Episodic Future Thinking Not Attributable to Deficits in Narrative Construction. The Journal of Neuroscience 31, 10262–10269.

Rolls, E.T., and Wirth, S. (2018). Spatial representations in the primate hippocampus, and their functions in memory and navigation. Prog Neurobiol 171, 90–113.

Saleem, A.B., Diamanti, E.M., Fournier, J., Harris, K.D., and Carandini, M. (2018). Coherent encoding of subjective spatial position in visual cortex and hippocampus. Nature 562, 124–127.

Silson, E.H., Chan, A.W.-Y., Reynolds, R.C., Kravitz, D.J., and Baker, C.I. (2015). A retinotopic basis for the division of high-level scene processing between lateral and ventral human occipitotemporal cortex. Journal of Neuroscience 35, 11921–11935.

Silson, E.H., Steel, A.D., and Baker, C.I. (2016). Scene-selectivity and retinotopy in medial parietal cortex. Frontiers in human neuroscience 10, 412.

Silver, M. A., & Kastner, S. (2009). Topographic maps in human frontal and parietal cortex. Trends in cognitive sciences, 13(11), 488–495.

Stachenfeld, K.L., Botvinick, M.M., and Gershman, S.J. (2017). The hippocampus as a predictive map. Nature neuroscience 20, 1643.

Suzuki, W.A. (2009). Perception and the Medial Temporal Lobe: Evaluating the Current Evidence. Neuron 61, 657–666.

Swisher, J. D., Halko, M. A., Merabet, L. B., McMains, S. A., & Somers, D. C. (2007). Visual topography of human intraparietal sulcus. Journal of Neuroscience, 27(20), 5326–5337.

Szinte, M., & Knapen, T. (2020). Visual Organization of the Default Network. Cerebral Cortex, 30(6), 3518–3527.

Turk-Browne, N. B. (2019). The hippocampus as a visual area organized by space and time: A spatiotemporal similarity hypothesis. Vision research, 165, 123–130.

Van Es, D. M., Van Der Zwaag, W., & Knapen, T. (2019). Topographic maps of visual space in the human cerebellum. Current Biology, 29(10), 1689–1694.

Wandell, B. A., Dumoulin, S. O., & Brewer, A. A. (2007). Visual field maps in human cortex. Neuron, 56(2), 366–383.

Wilming, N., König, P., König, S., & Buffalo, E. A. (2018). Entorhinal cortex receptive fields are modulated by spatial attention, even without movement. Elife, 7, e31745.

Wilson, C. L., Babb, T. L., Halgren, E., & Crandall, P. H. (1983). Visual receptive fields and response properties of neurons in human temporal lobe and visual pathways. Brain, 106(2), 473–502.

Zeidman, P., and Maguire, E.A. (2016). Anterior hippocampus: the anatomy of perception, imagination and episodic memory. Nature Reviews Neuroscience 17, 1

